# Simultaneous MRI-EEG during a motor imagery neurofeedback task: an open access brain imaging dataset for multi-modal data integration

**DOI:** 10.1101/862375

**Authors:** Giulia Lioi, Claire Cury, Lorraine Perronnet, Marsel Mano, Elise Bannier, Anatole Lécuyer, Christian Barillot

**Author notes:** Corresponding author: Giulia Lioi (,).

## Abstract

Combining EEG and fMRI allows for integration of fine spatial and accurate temporal resolution yet presents numerous challenges, noticeably if performed in real-time to implement a Neurofeedback (NF) loop. Here we describe a multimodal dataset of EEG and fMRI acquired simultaneously during a motor imagery NF task, supplemented with MRI structural data. The study involved 30 healthy volunteers undergoing five training sessions. We showed the potential and merit of simultaneous EEG-fMRI NF in previous work. Here we illustrate the type of information that can be extracted from this dataset and show its potential use. Our group is the second in the world to have integrated EEG and fMRI for NF, therefore this dataset is unique of its kind. We believe that it will be a valuable tool to

1. Advance and test methodologies to integrate complementary neuroimaging techniques (design and validation of methods of multi-modal data integration at various scales)
2. Improve the quality of Neurofeedback provided
3. Improve methodologies for de-noising EEG acquired under MRI
4. Investigate the neuromarkers of motor-imagery using multi-modal information

## Background & Summary

NF consists in providing real-time information to a subject about his own brain activity in order to train self-regulation of a specific brain function and is a promising brain rehabilitation technique for psychiatric disorders ^1^, stroke ^2,3^ and other neurological pathologies ^4^. NF approaches are usually based on real-time measures of brain activity using a single imaging technique, with the majority of applications relying on EEG and some recent ones employing functional imaging ^5,6^.

Recent studies ^7,8^ have shown the potential of combining different imaging techniques to achieve a more specific self-regulation. In particular, the combination of two complementary modalities such us fMRI and EEG seems particularly promising. fMRI offers fine spatial resolution (~mm) ^9^ but, as it is an indirect measure of the hemodynamic response, it has slow dynamics (~ s). On the other hand, EEG provides an excellent temporal resolution (~ms) and is a direct measure of neuronal activity. Its spatial resolution is however limited (~cm) by the volume conduction of cortical currents through the head tissues ^10^. The advantages of these imaging modalities are highly complementary and their integration is promising in applications requiring high temporal and spatial resolution such as NF training of specific brain areas.

More generally, the joint analysis of different imaging modalities can shed light on the complex link between anatomical, functional and electrophysiological properties of the brain ^11^. Integration of EEG and fMRI allows for an “augmented” analysis of the substrate neuronal dynamics, as the single modalities provide a partial estimation of the underlying neural activity. Joint EEG-fMRI analyses falls in two categories: asymmetrical and symmetrical ^12^. In the first information extracted from one methodology are integrated or drive the analysis of the second. In fMRI-driven EEG estimation for instance, fMRI activations are used as a prior to estimate putative EEG sources. Symmetrical approaches allowing for data fusion ^12^ have also been proposed, where a joint generative model is used. These approaches have been poorly explored, complexity and limited knowledge of neurovascular coupling being their main limitation. Similarly, in brain connectivity analysis, very few works attempted to integrate electrophysiological and hemodynamic information ^13^.

Collecting EEG in the ‘harsh’ environment of MR scanners entails however a series of technical challenges, primarily devoted to the reduction of the strong EEG artefacts generated by the rapidly varying MR gradient currents. The degradation of EEG SNR probably represents the main issue of simultaneous EEG-fMRI recordings and several de-noising solutions have been developed ^14,15^. However, so far, the corresponding datasets have rarely been released for use by other groups. Moreover, for the implementation of NF, simultaneous EEG-fMRI acquisition, artefact correction and processing need to be executed in real-time, adding complexity and challenge to hardware and software solutions.

To the best of our knowledge, simultaneous EEG-fMRI in a NF loop was implemented only by another research group to train emotional self-regulation ^7^: the dataset we share and describe here is therefore unique of its kind. It consist of 64-channels EEG and fMRI simultaneously acquired during a motor imagery NF task, complemented by structural MRI scans. Recordings were performed in two studies. In the first (XP1, Figure 1), 10 subjects performed bimodal and unimodal NF. The second study (XP2, Figure 2) involved 20 subjects performing bimodal NF using a bi-dimensional or mono-dimensional visual feedback.

**Figure 1.**
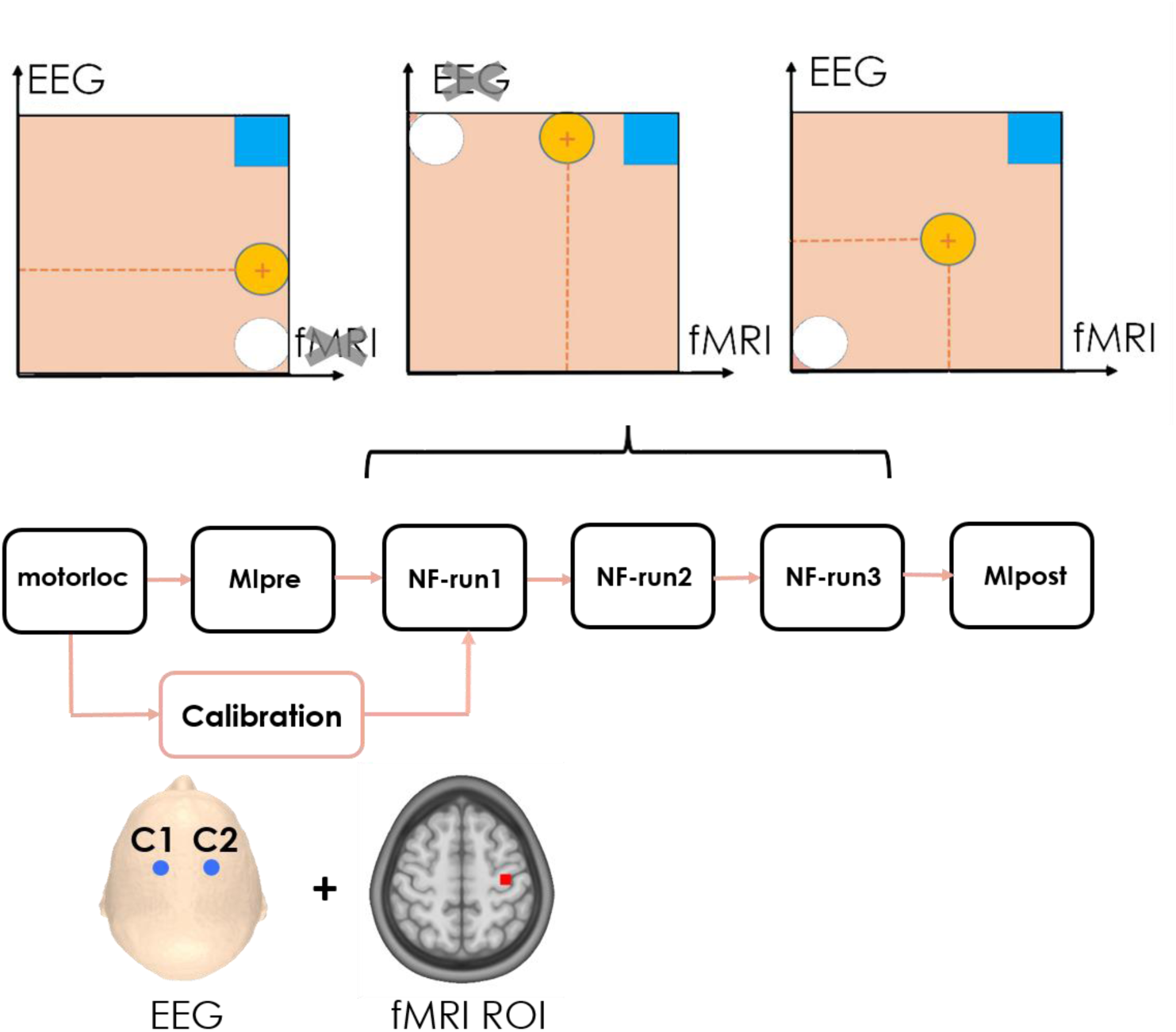
Experimental Protocol XP1.

**Figure 2.**
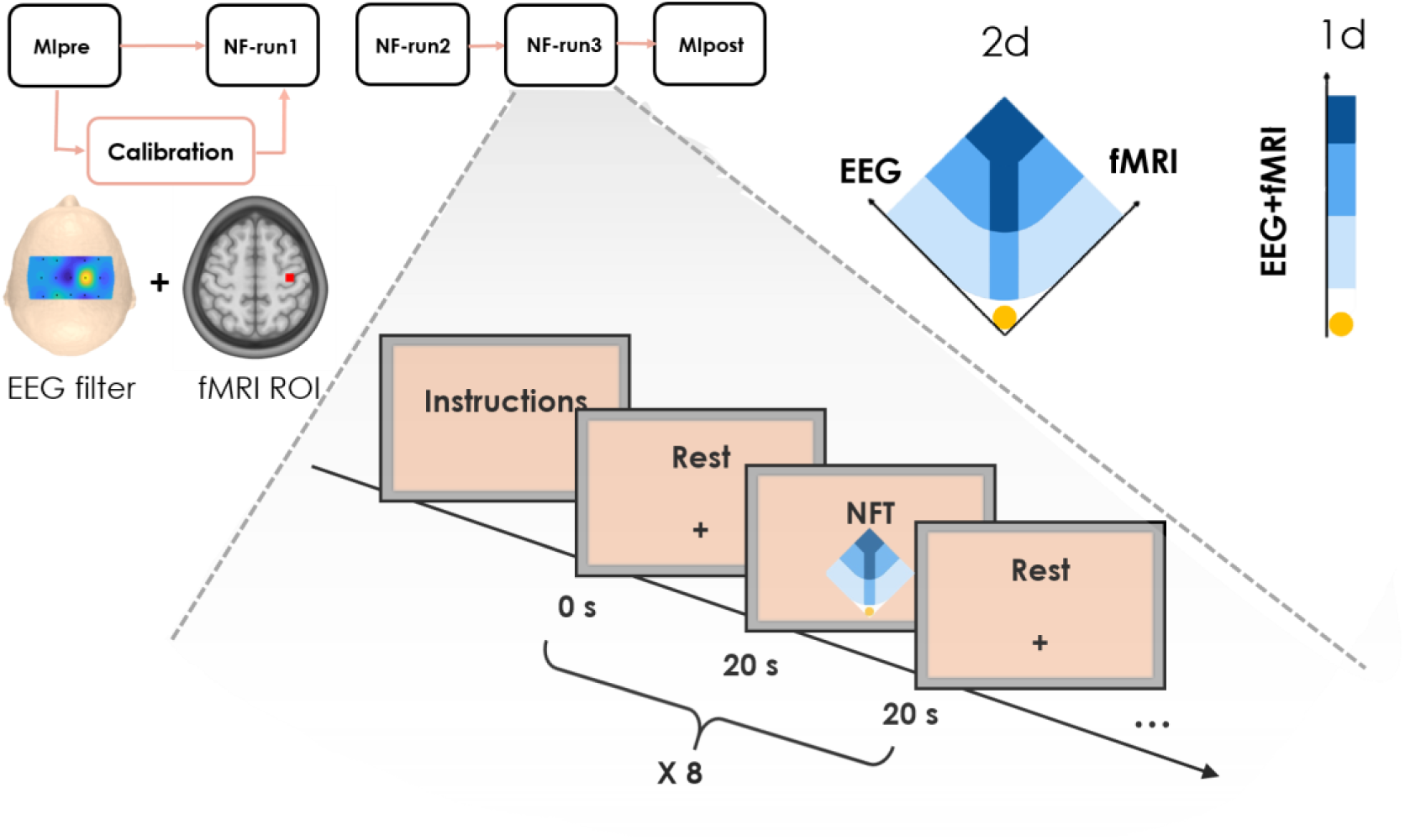
Experimental protocol XP2.

This dataset has potential to shed light on the coupling model underlying the EEG and fMRI signals, to advance methodologies for multimodal data integration and fusion techniques and test EEG de-noising algorithms. Since data were acquired during the execution of a simple task for whom patterns of neural pathways and activations are well known (motor imagery of the right hand), this dataset constitutes a simple model to develop and validate methods of data integration at various scales (activation maps, connectome).

This dataset is also of great interest for the NF research field, as it gives new perspectives to improve the estimation of NF scores, or the extraction of features of interest. We complemented this dataset with EEG and fMRI NF scores computed for each training session. These bi-modal NF data can be used to learn fMRI informed EEG in order to improve the quality of EEG only-NF ^16^. The development of such machine learning techniques has potential to increase the number and quality of NF training sessions and therefore advance clinical applications.

## Methods

### Participants

All experiments were performed according to the Helsinki declaration and under approval by the Institutional Review Board. Thirty right-handed NF-naïve healthy subjects were involved, of which 12 were female (mean age 33.7 ± 9.9 years old). All participants signed an informed consent after having been informed about the experimental procedure. Only 26 of them signed the consent to public their anonymized data, and were therefore included in the public datasets (Data Citation 1 and Data Citation 2).

### Data acquisition

The multimodal imaging NF platform used to acquire simultaneously EEG and fMRI data and to provide real-time NF has been described in ^17^ and a schematic is showed in Figure 3. It guarantees the acquisition, processing, NF features computation and visualization in real-time of the two data streams (EEG and fMRI). A sophisticated system of callbacks ensures full synchronization of the acquisition, stimuli and NF presentation processes. The platform includes a 64-channels MR-compatible EEG system (Brain Products GmbH, Gilching, Germany) and a 3T Verio MRI running VB17 (Siemens Healthineers, Erlangen), equipped with a 12 channels receiver head coil.

**Figure 3.**
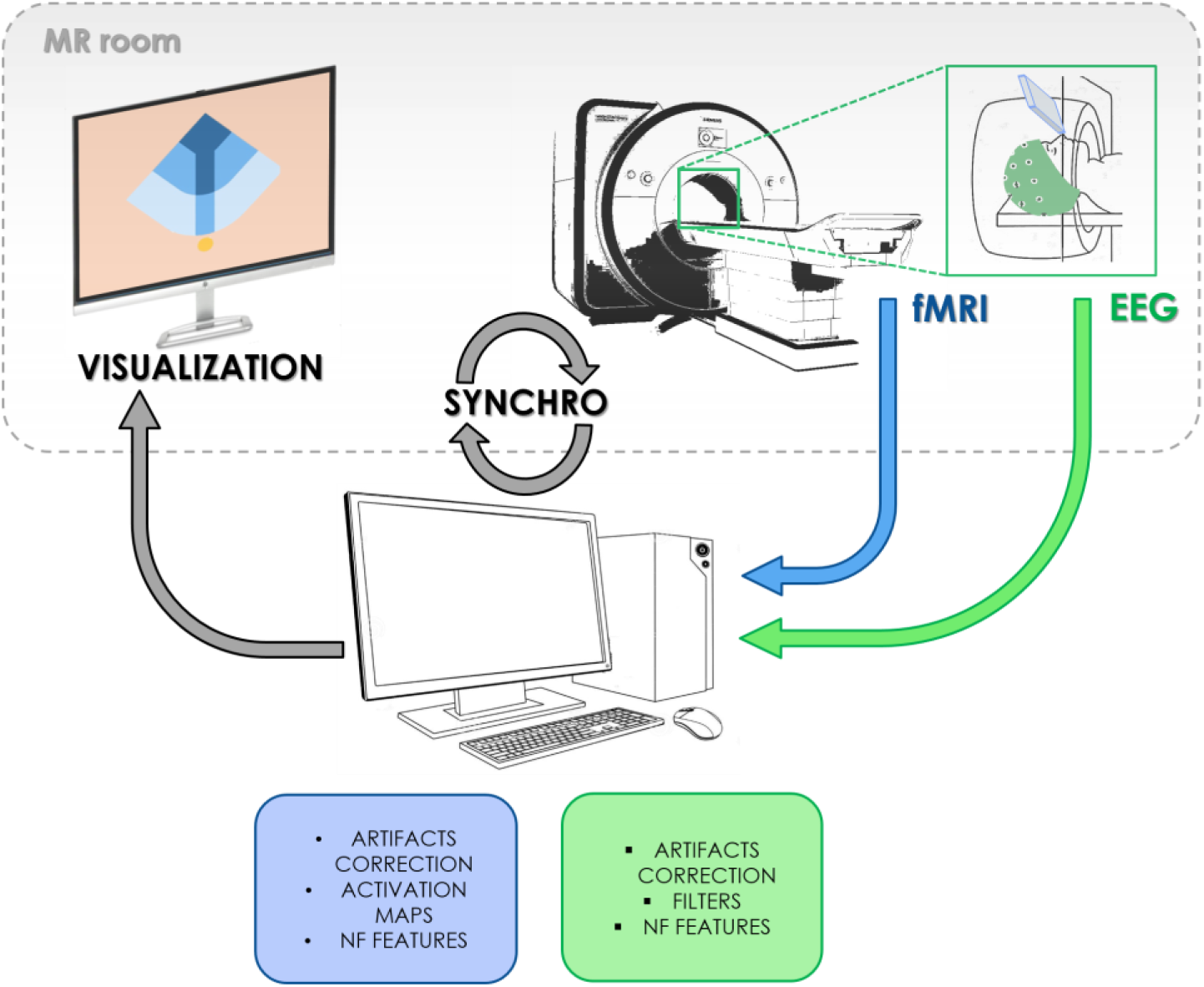
Schematic visualisation of the bimodal EEG-fMRI NF platform.

In line with the EEG vendor guidelines, a great care was put in the installation of the 64-channels EEG cap and reduction of electrode impedances. Electrodes impedances values estimated at the beginning of each run can be found in the header of the respective EEG file, as explained in the Data Records session. The subjects was then installed in the MR tube and electrodes impedances verified again after installation. Foam pads were applied to minimize head motion and for the whole duration of the experiment the subjects were asked to lie still in the MR bore. A LCD screen (NordicNeurolab Solutions, Bergen, Norway) was placed at the back of the MR scanner, a rear-facing mirror fixed on the top of the head coil allowed the subjects to view the screen.

For both studies, EEG recordings were sampled at 5 kHz with the reference electrode in FCz and ground in AFz. A 1mm isotropic 3D T1 MPRAGE structural scan was acquired. fMRI acquisition was performed by means of echo-planar imaging (EPI) whose settings changed from the first to the second study, as the repetition time (TR) could be further decreased thus improving the BOLD NF updating rate. Details about EEG and MRI data can be found in the Data Records session.

### Experimental Paradigm

#### XP1 (Data Citation 1)

The experimental paradigm of the first study has been described in detail in previous work ^8^ and is illustrated in Figure 1. Subjects were briefed about the task and familiarized with the NF metaphor (a ball moving in one or two dimensions depending on their brain activity). They were instructed to perform a kinaesthetic motor imagery of the right hand and to find their own strategy to control and bring the ball to the target. Specifically, they were informed that the NF would be a measure of laterality and that they had to maximize the activity only over the right hand motor region, therefore avoiding to image movements involving the two hands. Instructions were iterated verbally and on the screen at the beginning of NF run, together with the information about the direction (vertical, horizontal or both) along which the ball could be moved (Figure 1a). Participants were asked to hold still inside the MR bore and video monitoring allowed checking for movements of the subjects.

The experiment included six runs of a block-design alternating rest (20 s) and task (20 s). The first was a motor localizer (motorloc) where the subject was instructed to clench his right-hand every second for 5 min and 20s (8 blocks, starting with rest). A preliminary MI (MIpre) without feedback was then executed for 3 min and 20 s (5 blocks). Three NF blocks followed where the subject performed EEG-NF only (eegNF), fMRI-NF (fmriNF) and EEG-fMRI-NF (eegfmriNF) in random order across subjects. Each of the NF training run consisted of 10 blocks (6 min and 40 s). During NF runs the screen displayed a white ball moving in the vertical (condition eegNF, EEG laterality depicted the ball ordinate), or horizontal (condition fmriNF, BOLD laterality depicted the ball abscissa) or both dimensions (condition eegfmriNF) and a square representing the target. Finally, a transfer block of MI without NF (MIpost) was performed (3 min and 20 s), with the rationale of assessing the NF learning effect with respect to MIpre.

#### XP2 (Data Citation 2)

Methods and protocol for the second study have been described in ^18^ and are schematically showed in Figure 2. As for the first study, the experimental procedure and the NF task were carefully explained to participants before and during the experiment, in the form of written and verbal instructions. Similar instructions for kinaesthetic MI were given and the two NF metaphors (bi-dimensional 2d and unidimensional 1d) described. For the first group (1d) a ball moving on a mono-dimensional gauge was showed, and participants instructed to bring the ball towards the target (top, darkest areas of the gauge) by imaging to move their left hand. In the second group (2d) subjects could regulate separately the contribution of EEG and fMRI activities on the left and right axes, respectively (Figure 2).

Firstly a Mipre run (without feedback) was performed (5 min 20 s). Data collected from this session were used to identify the BOLD ROI and the optimal EEG filter (see next section). Three NF runs followed, each of 5 min and 20 s and spaced by a 1 min break. A MIpost block without feedback concluded the session.

### Real-Time Data processing and NF computation

During simultaneous EEG-fMRI acquisition, two main types of artefacts critically compromise the quality of EEG recordings: the rapid switching of the magnetic field gradients generate the so-called “gradient artefact”, while the heartbeat-induced movements of EEG electrodes generate the ballistocardiogram (BCG) artefact. Real-time correction of EEG artefact was performed using the Brain Vision Recview software. An average artefact subtraction approach ^14^ was used for gradient artefact correction, with four artefact templates. EEG data were then down-sampled to 200 Hz and filtered at 50 Hz (48 dB slope) for further processing. BCG artefact correction was then performed ^15^ using a moving template matching method with the following parameters: pulse period 800ms, correlation threshold 0.7 and amplitude ratio between the period examined and the pulse model ranging from 0.6 td 1.2. The moving pulse template was built averaging the 10 previous detected pulsed. The corrected EEG data were then sent to the NF control unit for feature extraction via a fieldtrip (https://github.com/fieldtrip/fieldtrip/) buffer solution.

Real-time fMRI pre-processing (slice-time and motion correction) was performed directly by the NF control unit using a custom Matlab script based on SPM 8 (FIL, Wellcome Trust Centre for Neuroimaging, UCL, London, UK).

#### XP1 (Data Citation 1)

Data collected during the motorloc session were used to identify a ROI for fMRI NF computation. fMRI scans were pre-processed for slice-time correction, spatial realignment and smoothing with SPM8 and activation maps were estimated with a GLM approach. A ROI over the left M1 was defined as the 9 × 9 × 3 voxels surrounding the maximum of activation. The right M1 ROI was defined as the symmetrical voxel across the mid sagittal plane. The activation from these two ROIs was included in the computation of the laterality NF feature and fed back to the subject during NFT. The EEG NF feature, on the other hand, was computed considering the EEG power in the (8-12) Hz band in the C1 and C2 electrodes (see the section NF features computation).

After pre-processing EEG features for the first study were computed according to the following expressions:

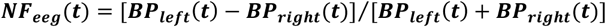

where ***BP***_***left***_(***t***) (***BP***_***right***_(***t***)) is the band power in the 8-12 Hz range at C1 (C2) during the NF task normalized by the power during rest. In other words. ***NF***_***eeg***_(***t***) is a measure of the motor imagery desynchronization laterality. The EEG laterality index was then smoothed,normalized and eventually translated as the ordinate of the NF ball and updated every 250 ms.

The fMRI laterality index was calculated on the corrected scans according to the expression:

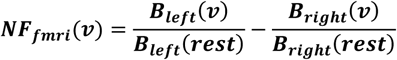

Here ***B***_***left***_(***v***) is the average of the BOLD signal in the left motor ROI identified during calibration at volume ***v*** normalized by the average signal in the same ROI over the last 6 volumes of the resting block ***B***_***left***_(***rest***). The BOLD laterality index was then smoothed and translated in the ordinate of the ball in the NF animation. The fMRI NF dimension was updated every 2 s (TR).

During rest blocks a cross was displayed and participants were asked to rest. During task, the screen displayed a cue (“move right”/“imagine right”) as well as the NF metaphor consisting in a ball moving on the horizontal (fmriNF), vertical (eegNF) or both axes (Figure 1) and target to reach.

#### XP2 (Data Citation 2)

For the second study, a more sophisticated calibration procedure was performed. Data collected during the MIpre session were used to identify an optimal spatial filter for EEG and a ROI based on the maximum of activation in the target motor area. As described in detail in ^18^, EEG data were pre-processed in Analyzer (Brain Product Software) and BCG artefacts were corrected with a semi-automated procedure. A Common Spatial Pattern (CSP) filter maximizing the difference in EEG power in the target band (8-30 Hz) between rest and MI blocks was estimated from a selection of motor channels. fMRI scans were pre-processed for slice-time correction, spatial realignment and smoothing with SPM8 and activation maps were estimated with a GLM approach. The selected ROI was defined as the voxel (9×9×3 mm) around the peak of activation in the target motor area (i.e. left motor cortex).

In XP2, EEG NF scores was a measure of event-related desynchronization (ERD) ^19^:

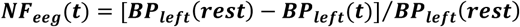

where ***BP***_***left***_(***t***) is the power in the 8-30 Hz frequency band and the ***BP***_***left***_(***rest***) is the average power over the resting block preceeding the NF training. ***NF***_***eeg***_(***t***) is a measure of the desynchronization occurring during motor imagery with respect to the baseline at rest. EEG NF scores were then smoothed over the last 4 values, normalized and converted in the visual feedback (ball movement) every 250 ms.

The fMRI feedback was calculated according to the following formula:

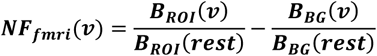

***B***_***ROI***_(***v***) is the BOLD signal in the motor ROI selected in the calibration step at volume ***v*** and it is divided by the corresponding signal averaged across the last 6 s of the previous rest block (to take in account the hemodynamic delay). ***B***_***BG***_(***v***) is the BOLD signal in a background deep slice, used here in order to normalize by global BOLD signal changes as recommended in ^20^. The fMRI NF index was averaged over the last three volumes and translated in a movement of the ball in the NF animation every second.

During NF runs the screen displayed instructions and the feedback which consisted of a ball moving in a two-dimensional plot for the 2d group or in a one-dimensional gauge for the 1d group (figure 1b).

### Code availability

A detailed description of bimodal EEG-fMRI NF platform is given in ^17^: the platform software package for real-time analysis and visualization is well documented but not publicly available. Python pipelines for the analysis of structural and functional MRI (Figure 8) are available on github (https://github.com/glioi/BIDS_fMRI_analysis_nipype), in form of commented jupyter notebooks. Other scripts used for the technical validation in this paper can be provided by the authors upon request.

## Data Records

The anonymized datasets are publicly available on the OpenNeuro repository in BIDS format ^21^. Even if the two datasets are very similar and, depending on the scientific purpose, can be jointly analysed, they were split in Dataset Citation 1 for XP1 data and Data Citation 2 for XP2 for the sake of clarity.

### XP1 (Data Citation 1)

The resource contains data from 10 subjects (18 GB), Table 1. The folder layout follows (both for MRI and EEG data) the BIDS specification folder hierarchy (Figure 4 A.).

**Table 1.** Participant demographic for XP1 (Data Citation 1): participants ID, age and sex is indicated, together with the NF tasks listed in the order they performed them.

| Data Citation 1 |  |  |  |
| --- | --- | --- | --- |
| participant_id | age | sex | feedback_task |
| sub-xp101 | 25 | M | fmri-nf, eeg-nf, eegfmri-nf |
| sub-xp102 | 27 | M | eeg-nf, eegfmri-nf, fmri-nf |
| sub-xp103 | 25 | M | fmri-nf, eeg-nf, eegfmri-nf |
| sub-xp104 | 31 | M | fmri-nf, eegfmri-nf, eeg-nf |
| sub-xp105 | 39 | M | eegfmri-nf, fmri-nf, eeg-nf |
| sub-xp106 | 36 | F | eegfmri-nf, eeg-nf, fmri-nf |
| sub-xp107 | 19 | M | eeg-nf, fmri-nf, eegfmri-nf |
| sub-xp108 | 29 | M | eeg-nf, fmri-nf, eegfmri-nf |
| sub-xp109 | 27 | F | fmri-nf, eegfmri-nf, eeg-nf |
| sub-xp110 | 26 | M | eegfmri-nf, fmri-nf, eeg-nf |

**Figure 4.**
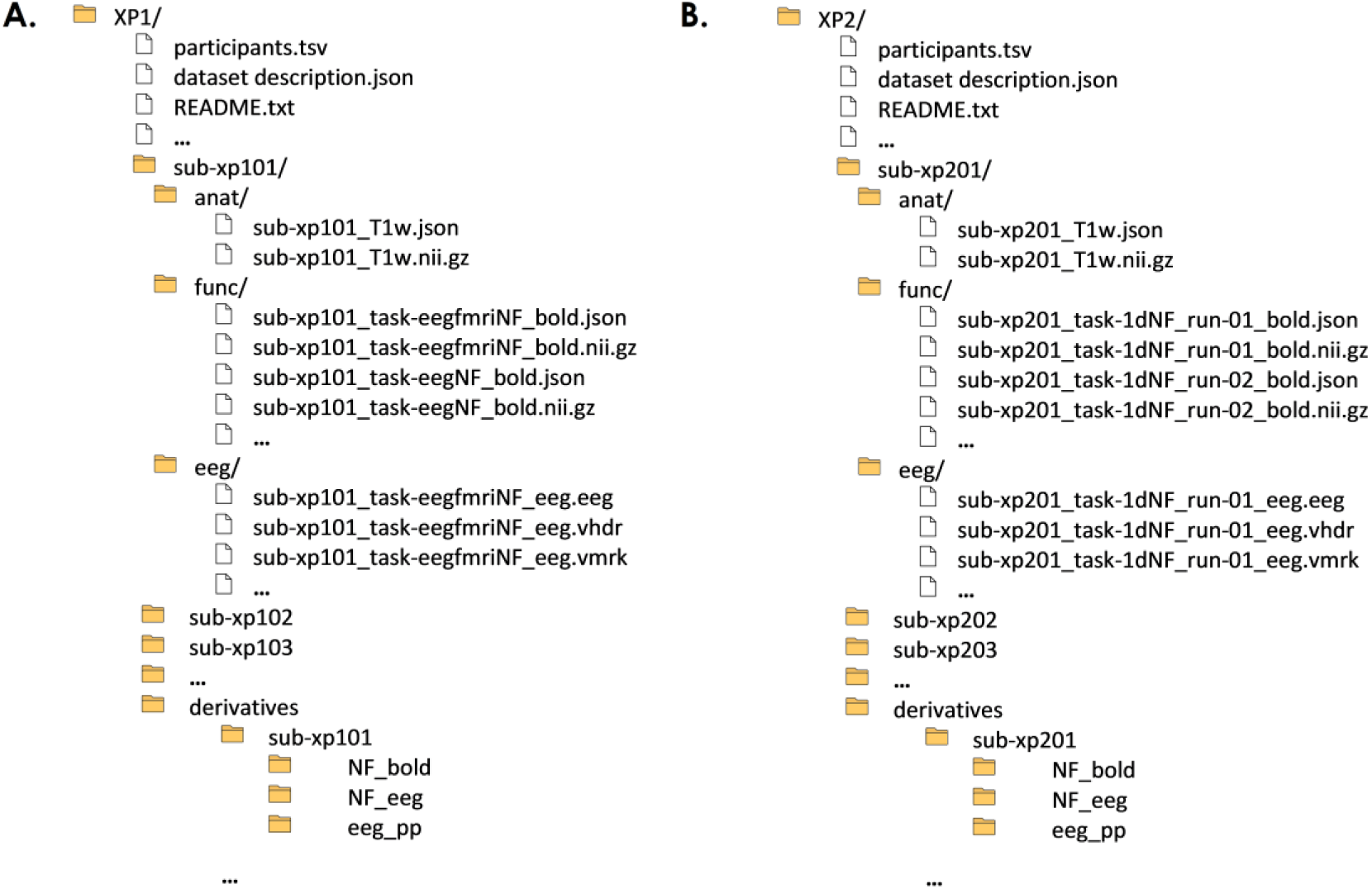
Folders architecture Data Citation 1 (A.) and Data Citation 2 (B.)

#### EEG DATA

##### Raw EEG

Raw EEG is sampled at 5kHz with FCz as the reference electrode and AFz as the ground electrode, and a resolution of 0.5 µV. Following the BIDS arborescence, raw EEG data for each task and subject can be found in

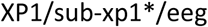

in Brain Vision Recorder format (File Version 1.0). Each raw EEG recording includes three files: the data file (*.eeg), the header file (*.vhdr) and the marker file (*.vmrk). The header file contains information about acquisition parameters and amplifier setup. For each electrode, the impedance at the beginning of the recording is also specified, giving therefore an additional indication of the quality of recording settings. For all subjects, channel 32 is the ECG channel. The 63 other channels are EEG channels. The marker file contains the list of markers assigned to the EEG recordings and their properties (marker type, marker ID and position in data points). Three type of markers are relevant for the EEG processing:

R128 (Response): is the fMRI volume marker to correct for the gradient artefact

S 99 (Stimulus): is the protocol marker indicating the start of the Rest block

S 2 (Stimulus): is the protocol marker indicating the start of the Task (that can be Motor Execution Motor Imagery or Neurofeedback). For technical reasons and for few runs, the first S99 marker was not correctly recorded: however it can be easily identified 20 s before the first S 2.

##### Preprocessed EEG

Following the BIDS structure, processed EEG data for each subject and task can be found in the pre-processed data folder:

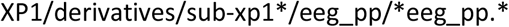

and following the Brain Analyzer format. Each processed EEG recording includes three files: the data file (*.dat), the header file (*.vhdr) and the marker file (*.vmrk), containing information similar to those described for raw data. In the header file of preprocessed data channels locations are also specified. In the marker file the location in data points of the identified heart pulse (R marker) are specified as well.

EEG data were pre-processed using BrainVision Analyzer II Software, with the following steps:

- Automatic gradient artefact correction using the artefact template subtraction method (sliding average calculation with 21 intervals for sliding average and all channels enabled for correction).
- Low Pass FIR Filter 50 Hz.
- Downsampling with factor 25 (200 Hz)
- BCG (pulse) artefact correction using a semiautomatic procedure (Pulse Template searched between 40 s and 240 s in the ECG channel with the following parameters: Coherence Trigger = 0.5, Minimal Amplitude = 0.5, Maximal Amplitude = 1.3. The identified pulses were marked with R).
- Segmentation relative to the first block marker (S 99) for all the length of the training protocol (last S 2 + 20 s).

##### EEG NF scores

Neurofeedback scores can be found in the. mat structures in

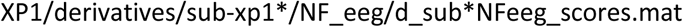

Structures names NF_eeg are composed of the following subfields:

- ID: Subject ID, for example sub-xp101
- lapC3_ERD: a 1×1280 vector of neurofeedback scores (4 scores per second)
- eeg: a 64×80200 matrix, with the pre-processed EEG signals according to the steps described above, filtered between 8 and 30 Hz.
- lapC3_bandpower_8Hz_30Hz: 1×1280 vector. Bandpower of the filtered signal with a laplacian centred on C3, used to estimate the lapC3_ERD.
- lapC3_filter: 1×64 vector. Laplacian filter centred on C3 channel.

#### MRI DATA

All DICOM files were converted to Nifti-1 and then in BIDS format (version 2.1.4) using the software dcm2niix (version v1.0.20190720 GVV7.4.0)

##### Raw functional MRI

Following the BIDS architecture (Figure 4 A.), the functional data and relative metadata are found for each subject in the following directory

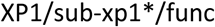

fMRI acquisitions were performed using echo-planar imaging (EPI) and covering the entire brain with the following parameters: TR=2 s, TE=23 ms, Resolution 2×2×4 mm^3^, FOV = 210×210mm^2^, Number of 4 mm slices: 32, No slice gap.

As specified in the relative task event files in XP1\ *events.tsv files onset, the scanner began the EPI pulse sequence two seconds prior to the start of the protocol (first rest block), so the first two TRs should be discarded. In task events files for the different tasks, each column represents:

- ‘onset’: onset time (sec) of an event
- ‘duration’: duration (sec) of the event
- ‘trial_type’: trial (block) type: rest or task (Rest, Task-ME, Task-MI, Task-NF)
- ‘stim_file’: image presented in a stimulus block: during Rest, Motor Imagery (Task-MI) or Motor execution (Task-ME) instructions were showed. On the other hand, during Neurofeedback blocks (Task-NF) the image presented was a ball moving in a square that the subject could control self-regulating his EEG and/or fMRI brain activity.

##### fMRI NF Scores

For each subject and NF session, a matlab structure with BOLD-NF features can be found in

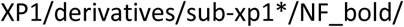

In view of BOLD-NF scores computation, fMRI data were preprocessed using SPM8 and with the following steps: slice-time correction, spatial realignment and coregistration with the anatomical scan, spatial smoothing with a 6 mm Gaussian kernel and normalization to the Montreal Neurological Institute (MNI) template

For each session, a first level general linear model analysis was then performed. The resulting activation maps (voxel-wise Family-Wise error corrected at p < 0.05) were used to define two ROIs (9×9×3 voxels) around the maximum of activation in the ipsilesional primary motor area (M1) and supplementary motor area (SMA) respectively.

The BOLD-NF scores were calculated as the difference between percentage signal change in the two ROIs (SMA and M1) and a large deep background region (slice 3 out of 16) whose activity is not correlated with the NF task. A smoothed version of the NF scores over the precedent three volumes was also computed.

The NF_bold structure is the following

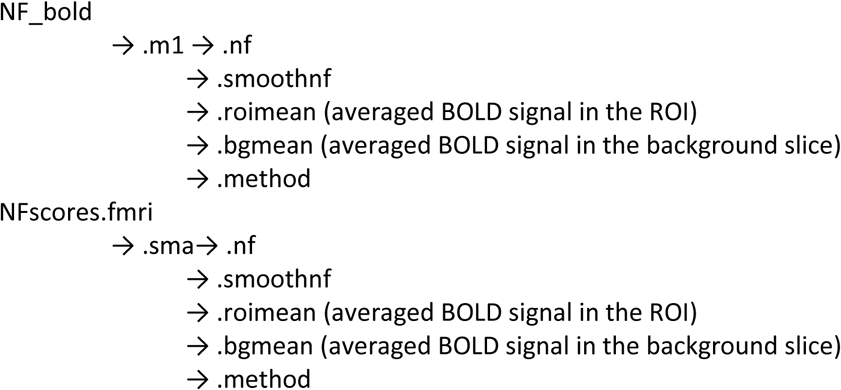

Where the subfield method contains information about the ROI size (.roisize), the background mask (.bgmask) and ROI mask (.roimask). More details about signal processing and NF calculation can be found in Perronnet et al. 2017 and Perronnet et al. 2018.

##### Anatomical MRI data

Following the BIDS standard, the functional data and relative metadata are found for each subject in the following directory

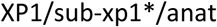

As a structural reference for the fMRI analysis, a high resolution 3D T1 MPRAGE sequence was acquired with the following parameters: TR=1900 ms, TE=2.26 ms, Resolution 1×1×1 mm^3^, FOV = 256×256 mm^2^, Number of slabs: 176.

Defacing of 3D T1 images was performed by the submitter using pydeface (https://github.com/poldracklab/pydeface)

### XP2 (Data Citation 2)

The resource contains data from 16 subjects (25 GB), as summarized in Table 2. Some subjects who participated to the study did not accept to make their data publicly available and were not included in the repository. The folder layout follows, both for MRI and EEG data, the BIDS format (Figure 5).

**Table 2.**
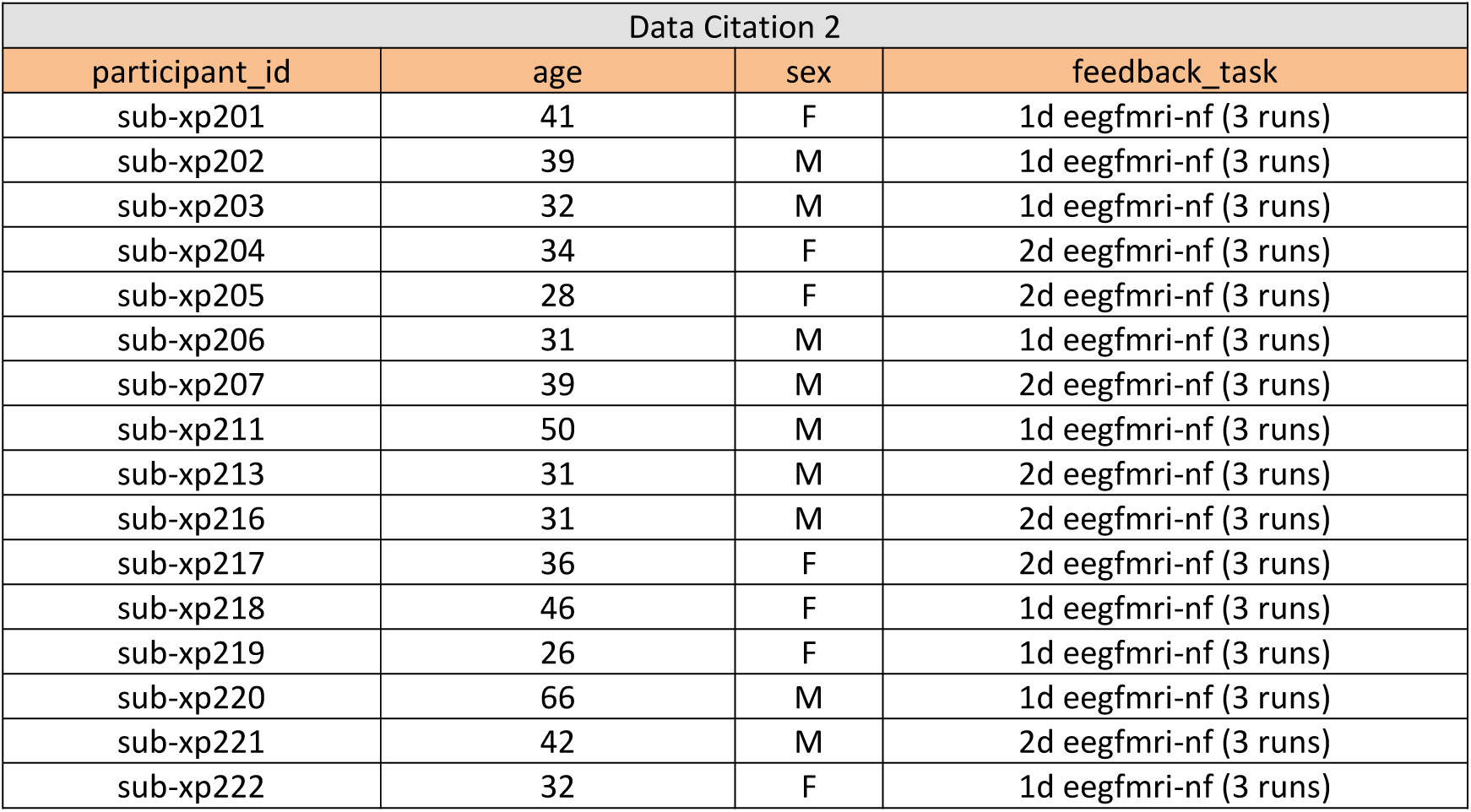
Participant demographic for XP2 (Data Citation 2): participants ID, age and sex is indicated, together with the NF tasks listed in the order they performed them.

| Data Citation 2 |  |  |  |
| --- | --- | --- | --- |
| participant_id | age | sex | feedback_task |
| sub-xp201 | 41 | F | 1d eegfmri-nf (3 runs) |
| sub-xp202 | 39 | M | 1d eegfmri-nf (3 runs) |
| sub-xp203 | 32 | M | 1d eegfmri-nf (3 runs) |
| sub-xp204 | 34 | F | 2d eegfmri-nf (3 runs) |
| sub-xp205 | 28 | F | 2d eegfmri-nf (3 runs) |
| sub-xp206 | 31 | M | 1d eegfmri-nf (3 runs) |
| sub-xp207 | 39 | M | 2d eegfmri-nf (3 runs) |
| sub-xp211 | 50 | M | 1d eegfmri-nf (3 runs) |
| sub-xp213 | 31 | M | 2d eegfmri-nf (3 runs) |
| sub-xp216 | 31 | M | 2d eegfmri-nf (3 runs) |
| sub-xp217 | 36 | F | 2d eegfmri-nf (3 runs) |
| sub-xp218 | 46 | F | 1d eegfmri-nf (3 runs) |
| sub-xp219 | 26 | F | 1d eegfmri-nf (3 runs) |
| sub-xp220 | 66 | M | 1d eegfmri-nf (3 runs) |
| sub-xp221 | 42 | M | 2d eegfmri-nf (3 runs) |
| sub-xp222 | 32 | F | 1d eegfmri-nf (3 runs) |

**Figure 5.**
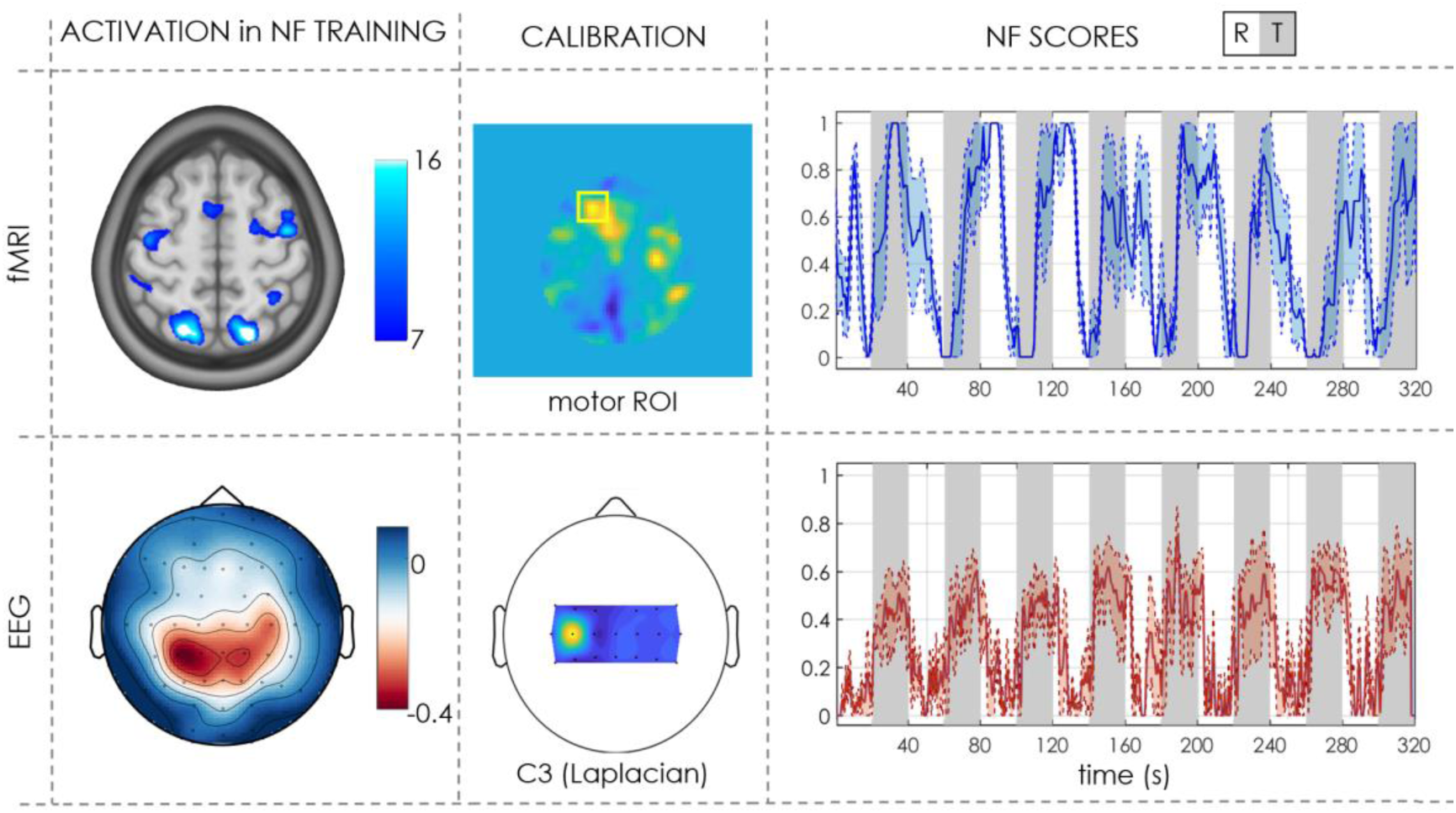
Example of fMRI and EEG activation and NF scores for one subject (Data Citation 2). The first column represents the activation (BOLD contrast for fMRI and ERD scalp distribution for EEG) during NF training. The second column show calibration features (ROI in the contralateral motor cortex for fMRI NF computation and Laplacian around the C3 electrode for EEG NF respectively). In the last column average (mean + std error) NF scores across 3 NF sessions are shown (rest blocks in white and NF task blocks in grey).

#### EEG DATA

EEG data were recorded, processed and saved as for Data Citation 1.

#### MRI DATA

All DICOM files were converted to Nifti-1 and then in BIDS format (version 2.1.4) using the software dcm2niix (version v1.0.20190720 GVV7.4.0)

##### Functional MRI

fMRI acquisition in XP2 slightly changed with respect to XP1. We used EPI covering the superior half of the brain with the following parameters: TR=1 s, TE=23 ms, Resolution 2×2×4 mm^3^, Number of 4 mm slices: 16, No slice gap. As for XP1 and as specified in the relative task event files in XP2\ *events.tsv, the scanner began the EPI pulse sequence two seconds prior to the start of the protocol (first rest block), so the first two TRs should be discarded.

In task events files for the different tasks, each column represents:

- ‘onset’: onset time (sec) of an event
- ‘duration’: duration (sec) of the event
- ‘trial_type’: trial (block) type: rest or task (Rest, Task-MI, Task-NF)
- ‘stim_file’: image presented in a stimulus block. During Rest or Motor Imagery (Task-MI) instructions were presented to the subject. During Neurofeedback blocks (Task-NF) the image presented was a ball moving in a square for the bidimensional NF (task-2dNF) or a ball moving along a gauge for the unidimensional NF (task-1dNF) that the subject could control self-regulating his EEG and fMRI brain activity.

fMRI preprocessing and NF scores computation was as for XP1.

##### Anatomical MRI data

Anatomical scans were acquired and defaced as for XP1.

## Technical Validation

The bimodal NF platform architecture for simultaneous EEG-fMRI recording is described in ^17^, together with details about the two layers data synchronization and preprocessing implemented in order to guarantee real-time NF presentation. Results relative to Data Citation 1 were published in ^8^, a study where the added value of bimodal NF with respect to the single modalities (EEG and fMRI only NF) was shown. Data Citation 2 experiment was intended to characterize a more subtle aspect: the impact of visual representation (2D or 1D) of bimodal NF on the NF performances ^18^. Finally, we are currently extending the bimodal NF paradigm to chronic stroke rehabilitation ^22^.

To illustrate the type of data and results we can extract from these datasets we show in Figure 5 an example of fMRI and EEG activation and NF scores for one subject (Data Citation 2). BOLD activation maps typically show activation of primary and supplementary motor areas and visual areas during the NF task as compared to rest. Similarly, the average ERD (8-30 Hz) scalp distribution for EEG presents a desynchronization over the motor electrodes, in particular in the contralateral region. BOLD NF scores show quite high (and consistent across sessions) activation of the selected ROI during the NF task, while EEG NF scores are lower and exhibit larger variability across sessions (which is to be expected given the higher NF update rate for EEG and the residual artefacts that affect EEG quality). The quality of EEG acquisition for each subject and session can also be verified in the corresponding header file where the impedance value for each electrode at the beginning of the recording is indicated: in all recordings the impedance was kept below 20 kΩ and in most cases values are below 10 kΩ.

To validate the data quality a classic analysis of EEG and fMRI patterns was performed over the 20 participants of Data Citation 2. EEG power spectrum was firstly estimated using a multitaper Hanning approach in the 8-30 Hz frequency band. ERDs for each block and NF run were then computed. The baseline for ERD computation was the 10 s interval before motor imagery execution (in order to exclude from the computation the event related synchronization ERS occurring at the end of the motor imagery task therefore in the first seconds of the following resting block). We performed a k-means cluster analysis on individual ERD and ERS features and identified two outliers (xp211, xp212) showing, at least in one NF run, artefactual ERD. These were excluded from further analysis of EEG patterns. Average ERD scalp distributions in the alpha (8-12 Hz) and beta (13-30 Hz) frequency bands were investigated, as well as temporal and frequency patterns, as shown in Figure 6. In line with results in literature, the average time-frequency map shows a desynchronization in the alpha band during the task block (and a less marked one in the beta band), together with a synchronization of the beta rhythm at the end of the motor task (i.e. in the first seconds of the rest block). Moreover ERD scalp distributions indicate that EEG activity involves mainly contralateral motor electrodes and is more specifically localized (C3, CP3) in the alpha band. We also performed EEG source reconstruction using an eLoreta approach and the template headmodel implemented in fieldtrip. A boundary element method (BEM) volume conduction model based on the template MRI (semiautomatically aligned with EEG fiducials) was computed and used for inverse model estimation. Average source ERD results (Figure 6c) indicate a desynchronization over the precentral end postcentral left gyri (corresponding to the sensory-motor cortex of the right upper limb).

**Figure 6.**
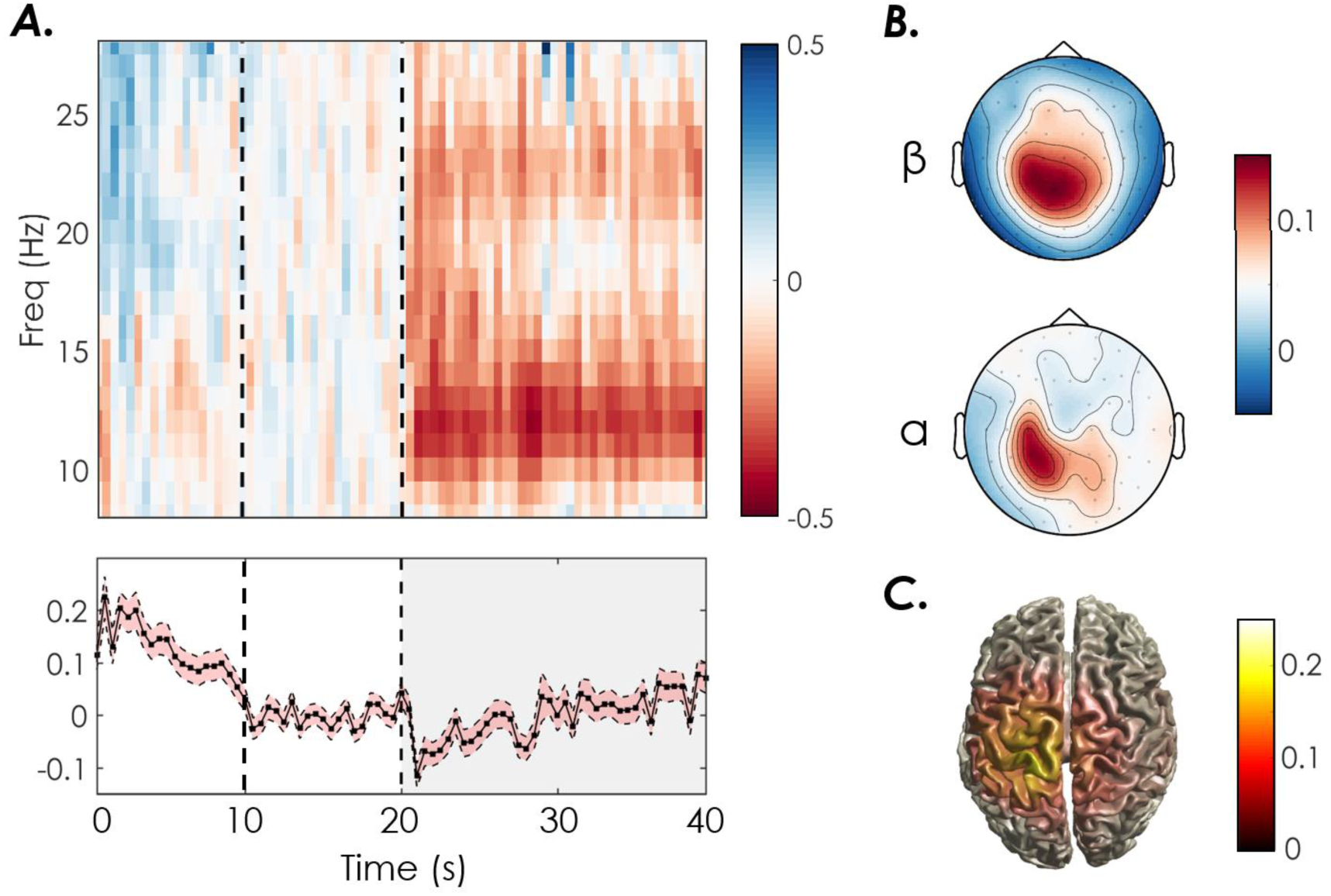
Average EEG ERD time-frequency patterns during NF training in XP2 (N=18 subjects XP2).A. Time-Frequency maps showing ERD (red) and ERS (blue) in the contralateral motor electrode and the corresponding ERD time series averaged across all frequencies (mean + standard error across subjects). B. ERD topographic maps in the alpha [8-12] Hz and beta [13-30] Hz frequency bands. C. ERD cortical maps.

fMRI analysis was performed using a processing pipeline based on Nipype, an open-source neuroimaging software that facilitates workflow reproducibility (https://github.com/nipy/nipype, version 1.2.0, Python 3.6). Preprocessing steps included slice timing and motion correction, detection of motion and intensity outliers, segmentation of anatomical image and coregistration, smoothing (fwhm=6). A fist-level and second-level GLM analysis were also performed across the three NF runs (canonical HRF, voxel-based analysis, Family-Wise error (FWE) correction p=0.05). Individual results were normalized to the Montreal Neurological Institute (MNI) template before performing a one sample T-test group analysis. More details about MRI data processing can be found in the processing scripts available on github (https://github.com/glioi/BIDS_fMRI_analysis_nipype) together with schematics of the workflow steps.

The motion-related and intensity outliers were detected using the ArtefactDetection tool implemented in Nipype (motion threshold=2, intensity z-score threshold=2): an average of 12.1 ± 5.3 scans per NF training run (corresponding to 3.3% of artefactual scans per session) were detected and excluded from the following GLM analysis. In line with results in literature, average BOLD activation maps (Figure 7) show significant activations in the premotor motor cortex and supplementary motor area (left and right), and in the left primary motor cortex during the motor NF task. These areas are involved in the planning and imagination of movement. The left posterior parietal cortex (PPC), that plays an important role in visuo-motor coordination, was also recruited during the motor imagery NF task, in line with influencing literature ^23^ indicating that PPC is generally active when feedback is presented visually. Results are quite robust across the two groups (2dNF and 1dNF) that exhibit largely overlapping contrast maps, as shown in Figure 7 B.

**Figure 7.**
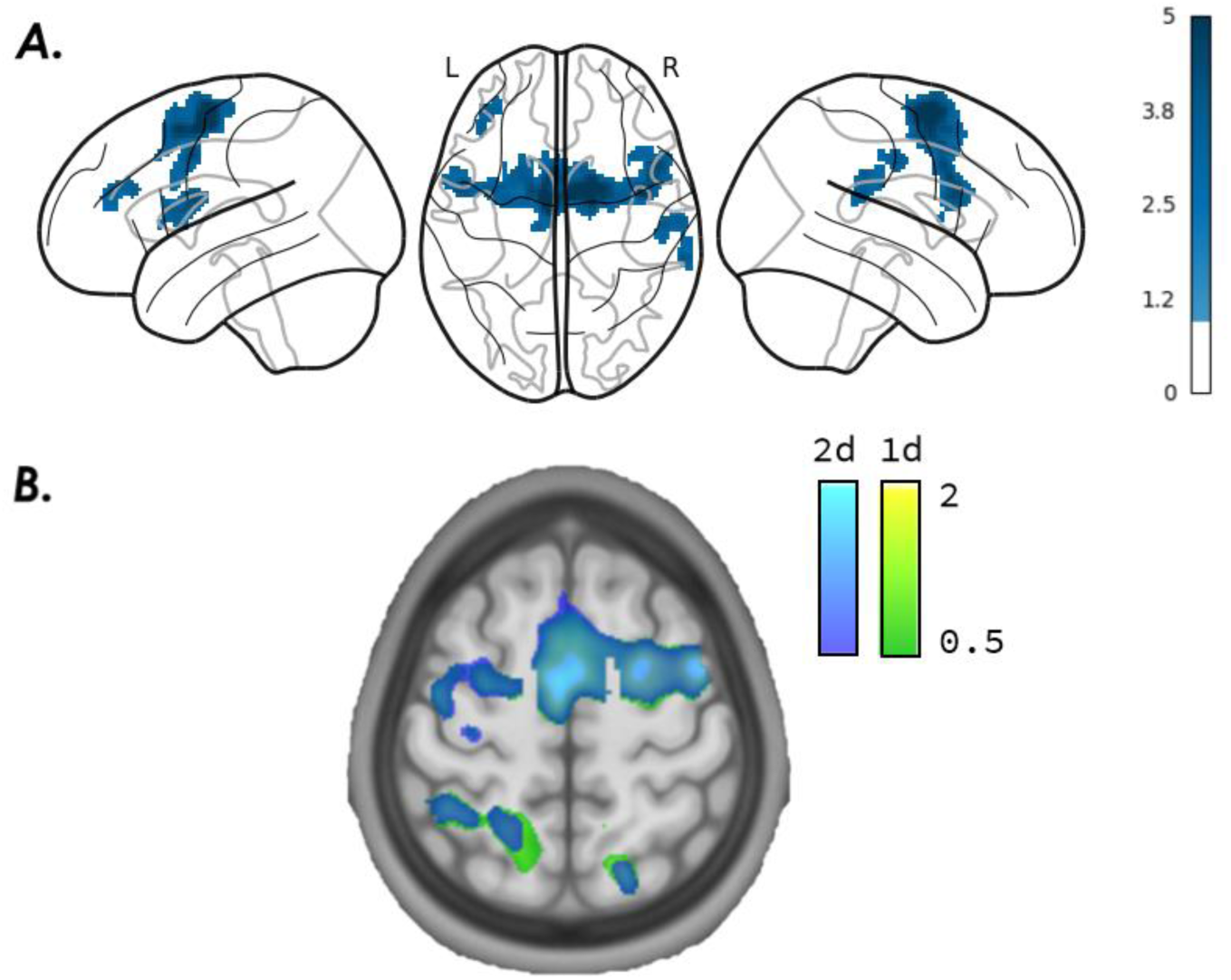
Group BOLD responses over three NF runs in XP2 (NF task>0, p=0.05 FWE corrected, voxel-based analysis). A. Average activation maps (N=20) thresholded at T-value=1. B. Contrast maps for the 2dNF group (blue) and 1dNF group (green).

To illustrate some possible uses for data fusion and integration, we performed two analysis, one on each dataset. The first one follows the methodology proposed in ^24^ to estimate sources localisation using a joint EEG and fMRI sparse model. The second shows the methodology to learn a fMRI informed EEG filter aiming at improving NF sessions using EEG only, as proposed in ^16^.

### 1. A data fusion approach for joint EEG-fMRI source estimation (XP1)

To estimate source location on the XP1 dataset, we applied the symmetric bi-modal method proposed by ^24^ to the motor localizer (motorloc) runs. The proposed approach uses a sparse regularisation for the sources and combines the two modalities EEG and fMRI to improve the spatio-temporal resolution of the sources estimation. The problem writes as follow:

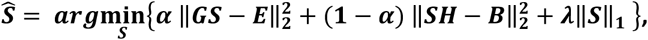

With ***S*** the estimated sources, ***G*** the leadfield matrix, ***E*** the observed EEG signals, ***B*** the observed BOLD signal and ***H*** a matrix linking EEG time samples to BOLD time samples using the hemodynamic response function. We chose different values for ***α*** (1 to consider EEG only, 0.6 the optimal balance between both modalities as found in ^24^ and 0 to consider fMRI only), and the optimal sparsity parameter *λ* = 2. We applied the model on all subjects having performed a motorloc run. Figure 8 shows the average results of source estimation from the joint EEG-fMRI model (***α*** = 0.6), from the EEG only model (***α*** = 1) and from fMRI only model (***α*** = 0). These results indicate that the combination of both modalities allows a more robust and complete localisation of the sources than when using only one modality, as even if fMRI provides a better spatial resolution, it can detect false positives, such as those outside motor areas.

**Figure 8.**
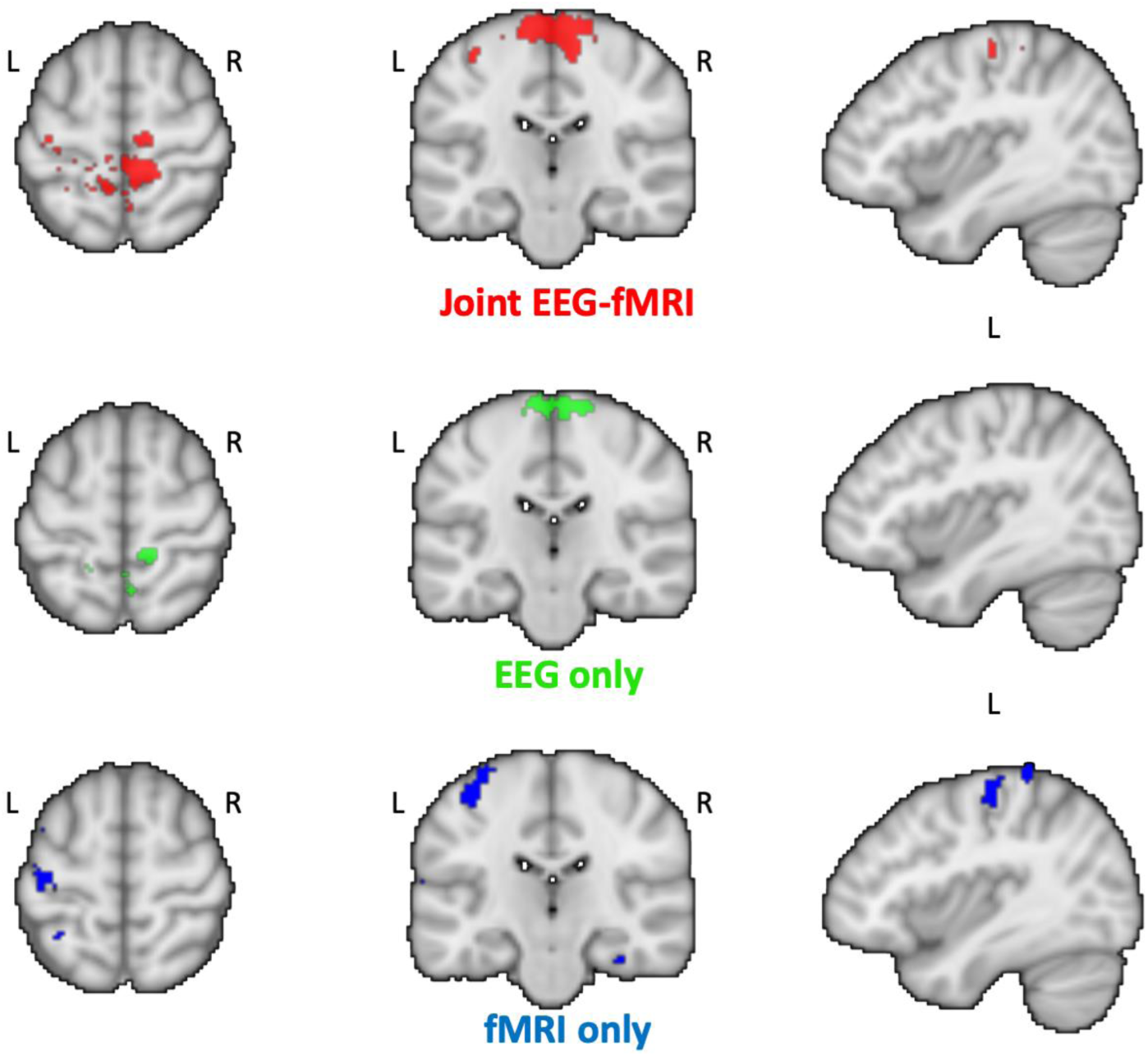
Joint EEG-fMRI source estimation average results. Source locations during motor execution (average across 8 subjects of XP1) estimated using the joint EEG-fMRI sparse model (red), EEG data only (α=1, green) or fMRI data only (α=0, blue).

### 2. A sparse EEG-informed fMRI model to improve EEG NF only (XP2)

We intended to learn fMRI-NF scores from EEG signal as proposed in ^16^ in order to determine if fMRI information could be integrated to EEG, on the dataset XP2. The learning and testing process is illustrated at Figure 9 panel A. The method proposes a model able to learn fMRI-NF scores from bi-modal NF sessions, using EEG signals only. The model learns some parameters of a design matrix composed by different frequency bands between 8 and 30 Hz, and different time delays to better match fMRI delays. A mixed norm is applied (*l*_1_ on the electrodes dimension and *l*_21_ on the frequency bands dimension) to the model to regulate the parameters. We excluded from the analysis the two subjects with a bad ERD, as detailed above. One of the sessions was used to learn the model, and the two other NF sessions are used to test the model. To validate the efficiency of the prediction, we correlated the estimated fMRI-NF scores with the fMRI-NF scores of the testing sessions, Figure 9 panel B shows boxplots in blue of such correlation across the dataset, for the learning phase and the testing phase. Information from fMRI could be extracted using EEG signals, as EEG signals could predict fMRI-NF scores with a median correlation of 0.34 with the ground truth. The model is quite well adapted to the data as shown by the learning step scores. Figure 9 also shows at panel B the correlation of the estimated EEG-fMRI-NF scores using only EEG-NF scores, with the ground truth that is the EEG-fMRI-NF scores. Figure 9 panel C shows an example of prediction on subject sub-xp216, the estimated NF scores have been learned on its first NF run and tested on its last run.

**Figure 9.**
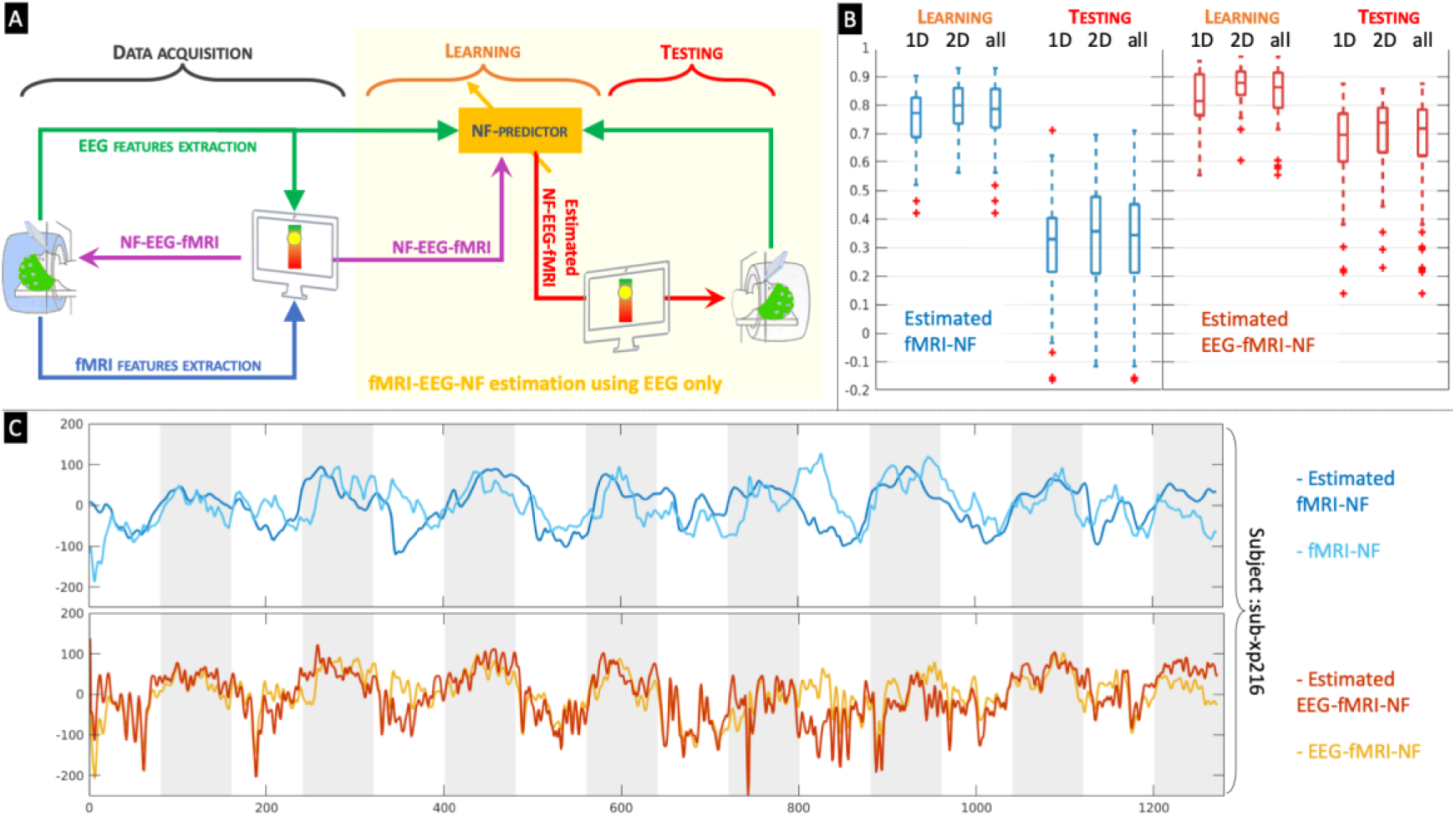
A sparse EEG-informed fMRI model to improve EEG NF only. Panel A: illustration of the method used to learn fMRI-NF scores from EEG signal. Panel B: Shows median and quartiles for correlations between estimated fMRI-NF scores and fMRI-NF scores for all subjects of the dataset XP2, for the learning step and the testing step. Results are also shown for the estimation of the bimodal EEG-fMRI-NF scores from EEG signal only. Panel C: Example for subject sub-xp216 run 3. The estimated scores were learned on its first NF run, light blue is the ground truth and dark blue the estimated fMRI-NF scores from EEG. Below, the yellow plot shows the bimodal NF score and in red its estimation using EEG only.

## Usage Notes

This dataset of simultaneously acquired EEG and fMRI during a NF motor imagery task is unique in literature and has potential to shed light on the coupling model underlying the EEG and fMRI signals, to advance methodologies for multimodal data integration and test EEG de-noising methods. The design of the platform and the data collection itself represented a challenge and a veritable research advancement: this constitute a major added value of this dataset but also implies that some technical challenges during data acquisition (especially for the first experiment XP1) led to missing training sessions in some subjects. Another limitation of this dataset is the quality EEG acquired during MRI imaging. As showed in the technical validation section and in previously published work, however, average results are in line with literature thus indicating the overall quality of EEG recordings. New (offline and real-time) EEG artefact correction algorithms may be efficiently tested on these data, as we provided raw and pre-processed EEG data. The efficacy of the proposed algorithms may for instance be assessed looking at quality of ERD patterns associated to the motor imagery (or execution) task.

Even if the definition of the BIDS standard for EEG is still evolving ^25^ we tried to adhere to this standard to have an easily and robustly exploitable EEG-MRI BIDS dataset, that can be processed using predefined and standardized pipeline (such as fMRIprep, BIDSHandler and various other BIDS-Apps, https://github.com/BIDS-Apps).

As mentioned in the previous section common and freely available software packages for EEG and MRI data processing were used to analyse data (SPM, Nipype, FieldTrip), except for EEG pre-processing for gradient and BCG artefact correction, for which the Brain Vision Analyser software (Brain Products GmbH, Gilching, Germany) was used. Other scripts used for the Technical Validation in this paper are provided by the authors as specified in the Code Availability session.

## Acknowledgements

MRI data acquisition was supported by the Neurinfo MRI research facility from the University of Rennes I. Neurinfo is granted by the the European Union (FEDER), the French State, the Brittany Council, Rennes Metropole, Inria, Inserm and the University Hospital of Rennes. The project was supported by the National Research Agency in the “Investing for 540 the Future” program under reference ANR-10-LABX-07-0, and by the “Fondation pour la Recherche Médicale” under the convention #DIC20161236427.

## Author contributions

G.L. wrote the manuscript, C.C contributed to the technical validation writing. L.P., C.B. and A.L. designed the studies. L.P. and M.M. collected the data. M.M and E.B. contributed to the design of the experimental platform. G.L. and C.C. conducted the data analysis, documented and standardized the dataset in BIDS format. All authors reviewed the manuscript.

## Competing interests

The authors declare no competing financial interests

## Notes

https://openneuro.org/datasets/ds002338/versions/1.0.1

https://openneuro.org/datasets/ds002336/versions/1.0.1

